# Sex-Biased Trajectories of Amygdalo-Hippocampal Morphology Change Over Human Development

**DOI:** 10.1101/661686

**Authors:** Ari M. Fish, Ajay Nadig, Jakob Seidlitz, Paul K. Reardon, Catherine Mankiw, Cassidy L. McDermott, Jonathan D. Blumenthal, Liv S. Clasen, Francois Lalonde, Jason P. Lerch, M. Mallar Chakravarty, Russell T. Shinohara, Armin Raznahan

## Abstract

The amygdala and hippocampus are two adjacent allocortical structures implicated in sex-biased and developmentally-emergent psychopathology. However, the spatiotemporal dynamics of amygdalo-hippocampal development remain poorly understood in healthy humans. The current study defined trajectories of volume and shape change for the amygdala and hippocampus by applying a multi-atlas segmentation pipeline (MAGeT-Brain) and semi-parametric mixed-effects spline modeling to 1,529 longitudinally-acquired structural MRI brain scans from a large, single-center cohort of 792 youth (403 males, 389 females) between the ages of 5 and 25 years old. We found that amygdala and hippocampus volumes both follow curvilinear and sexually dimorphic growth trajectories. These sex-biases were particularly striking in the amygdala: males showed a significantly later and slower adolescent deceleration in volume expansion (at age 20 years) than females (age 13 years). Shape analysis localized significant hot-spots of sex-biased anatomical development in sub-regional territories overlying rostral and caudal extremes of the CA1/2 in the hippocampus, and the centromedial nuclear group of the amygdala. In both sexes, principal components analysis revealed close integration of amygdala and hippocampus shape change along two main topographically-organized axes – low vs. high areal expansion, and early vs. late growth deceleration. These results bring greater resolution to our spatiotemporal understanding of amygdalo-hippocampal development in healthy males and females and discover focal sex-differences in the structural maturation of the brain components that may contribute to differences in behavior and psychopathology that emerge during adolescence.

**SIGNIFICANCE STATEMENT:** The amygdala and hippocampus are implicated in several developmentally-dynamic and sex-biased psychiatric disorders, but the spatiotemporal organization and sex-biased patterning of amygdalo-hippocampal maturation remains unclear in humans. Here, by integrating new methods for analysis of longitudinal neuroimaging data, we resolve the developmental milestones and spatial gradients that organize human amygdalo-hippocampal maturation. Each structure’s volume follows a tri-phasic, curvilinear growth trajectory which - for the amygdala - shows rapid male-female size divergence in mid-adolescence through delayed growth deceleration in males. Spatially fine-grained shape analyses localize these sex differences, and further reveal highly orchestrated shape changes across the amygdala and hippocampus that are organized by two topographical gradients. These data provide a new framework for understanding amygdalo-hippocampal organization in human development.

## INTRODUCTION

The amygdala and hippocampus are two adjacent allocortical structures that play key roles in learning, memory, and emotion (Burgess, Maguire, & O’Keefe, 2002; Maren, De Oca, & Fanselow, 1994; Scoville & Milner, 1957), and are also implicated in the neurobiology of diverse psychiatric and neurodevelopmental disorders including autism, attention deficit hyperactivity disorder, psychopathy, anxiety disorders, major depression, and schizophrenia (Haukvik, Tamnes, Söderman, & Agartz, 2018; Kalin, 2017; Shaw & Rabin, 2009; Yang et al., 2009). Although these domains of brain function and dysfunction all show marked developmental sex-differences between childhood and early adulthood (Rutter, Caspi, & Moffitt, 2003; Satterthwaite et al., 2015), relatively little is known regarding the spatiotemporal patterning of amygdalo-hippocampal development in males and females.

To date, there is strong evidence from longitudinal structural neuroimaging for robust effects of age, sex, and pubertal status on bulk volume of the amygdala and hippocampus (Goddings et al., 2014; Herting et al., 2018; Walhovd et al., 2005), but we lack similarly large-scale longitudinal studies of age and sex effects on regional amygdalar or hippocampal shape in typically-developing humans (Gogtay et al., 2006). The importance of securing spatially fine-grained analyses of amygdalo-hippocampal maturation is highlighted by recent evidence that the volume of different hippocampal subfields may follow distinct developmental tempos (Tamnes et al., 2018). Here, we extend the largest existent longitudinal study of subcortical morphology in human development (Raznahan, Shaw, et al., 2014) to chart structural maturation of the human amygdala and hippocampus at high spatiotemporal resolution between ages 5 and 25 years within a single-site cohort. Our analyses seek to advance knowledge of normative amygdalar and hippocampal development in two main directions.

First, we re-visit bulk volume development in the amygdala and hippocampus (Goddings et al., 2014; Gogtay et al., 2006; Tamnes et al., 2017, 2018), by (i) applying cutting-edge, multi-atlas algorithms for automated subcortical segmentation (Chakravarty et al., 2013; Pipitone et al., 2014) to a large, single-center longitudinal neuroimaging study of human brain development (Giedd et al., 2015), and (ii) estimating developmental trajectories with semi-parametric spline-based methods, which offer several advantages over classical polynomial approaches to curve-fitting (Mills & Tamnes, 2014). We combine these methodological advances to capture curvilinear trajectories of amygdala and hippocampal volume across the transition from childhood and early adulthood in each sex, and pinpoint key maturational markers (Fjell et al., 2010; Raznahan, Shaw, et al., 2014; Shaw et al., 2008) that have remained unresolved for the amygdala and hippocampus: age of fastest volume change, age of greatest growth deceleration, and age at attainment of peak volume.

Second, we move beyond analysis of bulk volume by modelling trajectories of surface area change at 5,245 points (vertices) across the amygdala and hippocampus to provide reference, surface-based “maturation maps” for these structures that are in line with those that are already available for the cortex, striatum, pallidum, and thalamus (Raznahan, Shaw, et al., 2014; Tamnes et al., 2017). Such maps are essential for (i) localizing “hot-spots” of developmentally-emergent sex differences in brain anatomy (Raznahan, Lee, et al., 2014), (ii) identifying spatial gradients in maturation tempos within individual structures (Raznahan, Shaw, et al., 2014), and (iii) identifying distributed patterns of coordinated maturation across structures (Alexander-Bloch, Giedd, & Bullmore, 2013). These surface-based approaches provide a valuable complement to classical segmentation-based approaches for analysis of sub-regional anatomy of the amygdala and hippocampus by recovering organizational gradients that vary within or cut across canonical subfields/subnuclei (Brunec et al., 2018; Collin, Milivojevic, & Doeller, 2015; Poppenk, Evensmoen, Moscovitch, & Nadel, 2013; Sah et al., 2003; Vos de Wael et al., 2018). Extending this notion, we pursue joint surface-based analysis of amygdalar and hippocampal anatomy, to allow detection of coordinated maturation between these structures – motivated by evidence for coordinated functional networks that overlap both structures (Ji et al., 2019; Phelps, 2004; Richardson, Strange, & Dolan, 2004).

## METHODS

### Participants

Our longitudinal study included 1,529 structural magnetic resonance imaging (sMRI) scans gathered from 792 healthy males and females (403 M/ 389 F) between 5 and 25 years of age (**Table 1**). The number of scans per person ranged from 1 to 6, with 51% of participants providing 2 or more scans (26% with 3 or more scans). Participant recruitment and assessment were conducted as previously detailed (Giedd et al., 1999). Briefly, participants were recruited for the study through local advertisements and exclusion criteria included: Full scale IQ less than 80 (as determined by age-appropriate Wechsler Intelligence Scale tests) and presence of any neurological disease or psychiatric illness (as determined by phone screen and completion of the Childhood Behavior Checklist questionnaire). The research protocol was approved by the Institutional Review Board at the National Institute of Mental Health, and written informed consent or assent was obtained from all children who participated in the study, as well as consent from their parents if the child was under the age of 18.

**Table 1.**
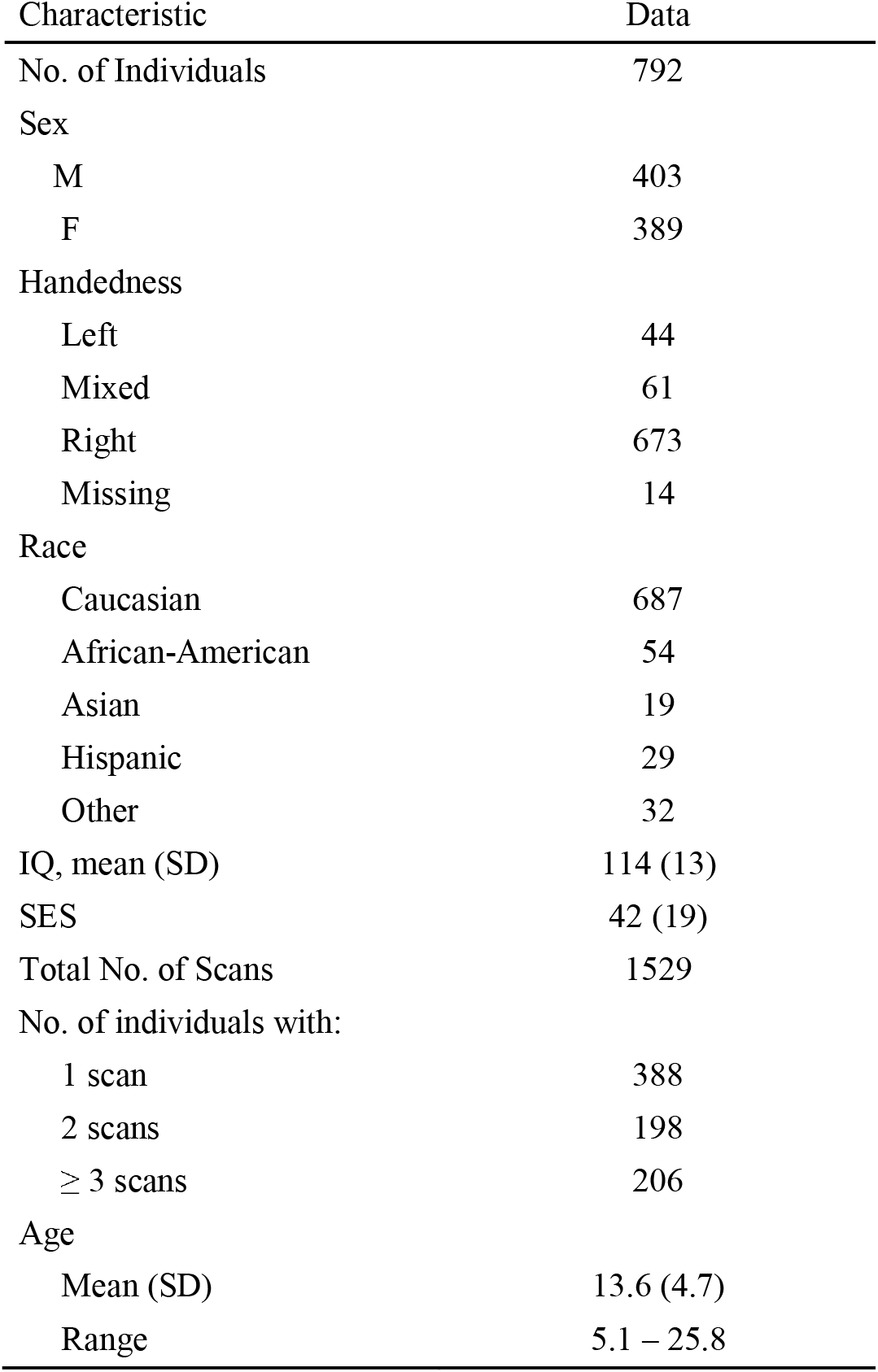
Participant Characteristics. SES, socioeconomic status as measured by the Hollingshead Scales

### Neuroimaging

All sMRI scans were obtained on the same 1.5-T General Electric Signa scanner located at the National Institutes of Health in Bethesda, MD. All scans were T-1 weighted images with contiguous 1.5-mm axial slices and 2.0-mm coronal slices. We used a 3D spoiled-gradient recalled-echo sequence with parameters as follows: 5 ms echo time, 24 ms repetition time, 45° flip angle, 256 × 192 acquisition matrix, 1 excitation, and 24 cm field of view. All scans met 2 quality assessment criteria: (i) absence of visible motion artifact in raw scans prior to pre-processing (Alexander-Bloch et al., 2016), and (ii) visual inspection of subcortical segmentation labels for each scan using montages of output post-processing by MAGeT brain (Raznahan, Shaw, et al., 2014).

### Subcortical Segmentation

Subcortical structures were automatically identified and segmented using the multi-atlas segmentation algorithm, MAGeT Brain (Chakravarty et al., 2013; Pipitone et al., 2014; Winterburn et al, 2013). Briefly, MAGeT Brain performs regional segmentation using *in vivo* atlases generated from high-resolution T1- and T2-weighted images. The atlases were generated from 2 males and 3 females. These atlases are customized to 21 randomly selected candidate subjects from the NIH Human Brain Development in Health Study (Giedd et al., 2015) and are then used as the templates to which all other scans are registered for segmentation. Thus, a study-specific library of 105 amygdala and hippocampus segmentations is produced for each scan (5 atlases × 21 templates), and conclusive segmentation is determined by using the label that most often occurs at a specific location (i.e., a voxel-wise majority vote). This method compares favorably in terms of reliability against previous gold-standard manual tracing definitions in a sample of adolescent brain images (Dice Kappa = 0.88; Pipitone et al, 2014). All scans then underwent a detailed quality control inspection (rater: P.K.R.) to identify gross segmentation errors and artifacts.

To determine the shape of the amygdala and hippocampus, MAGeT-Brain uses a marching cubes algorithm to create surface-based visualizations – which were then smoothed using the AMIRA software package (Visage Imaging) in a group-specific atlas of the original 5 atlas images (Voineskos et al., 2015). The nonlinear portions of the transformations were concatenated and averaged across all 21 input templates resulting in 105 possible surface representations per subject, which were then combined by estimating the median coordinate representation at each point. One-third of the adjoining triangle’s surface area was assigned to each vertex. All surface assignments from associated triangles were summed to produce the estimated surface area value at each vertex, and then these values were blurred using a surface-based diffusion-smoothing kernel (5 mm). These vertex-level surface area estimates at a total of 5,245 points (right amygdala [1,405], left amygdala [1,473], right hippocampus [1,215], and left hippocampus [1,152]) were used to spatially map amygdalo-hippocampal shape variation as a function of age and sex. Vertex-level area analyses also provided insights into regional drivers of global changes in amygdalar and hippocampal size given that the sum of vertex area for each structure was a close proxy for structure total volume (cross-scan correlations between total nuclear surface area and volume: r_AMY_=0.96,r_HIPP_=0.90).

### Experimental Design and Statistical Analysis

Data analysis and visualization was conducted within the R computing environment (R Core Team, 2015) using the following packages and their dependencies: nlme (Pinheiro et al., 2018), gamm4 (Wood & Scheipl, 2017), ggplot2 (Wickham, 2016), and dplyr (Wickham, Francois, Henry, & Muller, 2018). Age and sex effects on anatomical variation were estimated using generalized additive mixed models (GAMMs), which allow flexible and efficient estimation of developmental curves with several advantages over traditional polynomial regression models. These advantages include greater robustness of local curve estimation to variation in age range of scans included, and the ability to detect patterns of curvilinearity that do not fit within the classes offered by standard polynomials (Mills & Tamnes, 2014).

#### Testing for and characterizing sex differences in anatomical trajectory shape

Statistical inference regarding sex differences in the developmental trajectory of the bulk bilateral volume of each structure, and on the area of individual vertices within each structure, were based on p-values associated with the second smooth term in semiparametric model [1] below. Importantly, a nested random effect term was included to account for the dependency of scans within individuals, and individuals within families. For vertex-wise analyses, these p-values were corrected for multiple comparisons using the false discovery rate (FDR) method, with q (the expected proportion of falsely rejected nulls) set at 0.05 (Genovese, Lazar, & Nichols, 2002). Thus, sex differences in trajectory shape were assessed using the following model:

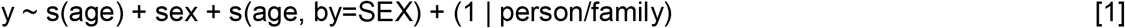

Where y represents the anatomical variable being modelled, s() is a fixed degrees of freedom thin plate regression spline with k (the number of knots) set at 5. This value of k was set to match the highest estimated degrees of freedom across both spline terms in [1] from fits of the model to bulk amygdala and hippocampal volumes using penalized smoothing. We chose fixed degree of freedom regression splines over penalized smoothing splines for inference regarding sex-differences in trajectory shape because this approach defined - for each anatomical metric of interest - a single, minimal degree of freedom for use in bootstrap analysis of maturational milestones in male and female developmental trajectories (see below).

Bootstrap methods were used to estimate the mean and 95% confidence intervals for three different developmental markers extracted from the sex-specific volume trajectories in each structure: age at attainment of peak volume, age at fastest volume change, and age at greatest negative inflection in volume change. Specifically, we generated 1000 bootstrap samples of longitudinal neuroimaging data by drawing the scans for 792 individuals (the number of unique individuals in our dataset) from our full cohort with replacement (using the *sample* function in R), and we then estimated sex-specific trajectories of volume for each bootstrap sample using model [1] above. We identified the age at attainment of peak structure volume for each of these sample-specific volume trajectories, and also calculated point rate of volume change trajectories to identify (i) the age at peak volume change, and (ii) the age at greatest negative inflection in rate of volume change (i.e., growth deceleration) for each sample. The age of fastest growth deceleration was defined as the age with the most negative point rate of change in the rate of volume change (i.e., second derivative or “growth deceleration”). The distributions of these values across all 1000 bootstrap samples were used to derive structure-specific means (μ_boot_) and 95% confidence intervals (CI_boot_) for all three developmental milestones. For each bootstrap resample, we computed the difference between male and female timing of each developmental milestone and used the resulting milestone difference distribution to test for statistical significance of sex differences. We concluded a significant sex-difference for a developmental milestone when more than 95% of bootstrap resamples resulted in same-direction sex difference in milestone timing (i.e., p_boot_ < 0.05).

#### Mapping spatial gradients of anatomical change within each sex

Within each sex, effects of age on the area of individual vertices were estimated using the following model:

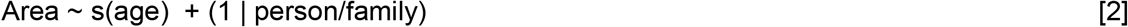

Where s() is a penalized regression spline term for age. We chose a penalized regression spline for this analysis (versus fixed degree of freedom regression splines as in model [1]) in order to achieve vertex-specific optimization of trajectory smoothness, given the focus of these analyses on regional variations in trajectory shape within each sex (rather than statistical inference regarding male-female differences). Regional variations in trajectory shape were visualized in each sex using two complementary approaches. To examine regional gradients in the overall magnitude of surface area change over development, we generated vertex-wise maps of percent area change across the 20-year span (ages 5 to 25 years) of our study for the hippocampus and amygdala. These maps provide a simple representation of inter-regional differences in the overall magnitude of anatomical change for each structure, but do not consider possible variations in the tempo of anatomical change between ages 5 and 25 years. We therefore also used the developmental trajectories for surface area provided by model [2] above to calculate annualized percent surface area change trajectories at each vertex (at 400 evenly-spaced age points across our study range), and then submitted these matrices (where rows are vertices, columns are ages, and values are point estimates of rate of area change) to Principal Components Analyses (PCA). The resulting PCs of area change could then be visualized on the amygdala and hippocampal surface to define dominant spatial variations in the temporal dynamics of amygdalo-hippocampal shape. Critically, PCA was conducted on a concatenated matrix of hippocampal and amygdala area change data, allowing us to define coordinated patterns of anatomical change that spanned both structures. We selected the optimal number of components as 2 - based on the scree plot for the resulting PCA model. Vertex loadings onto these components were visualized as separate surface maps for each component, as well as a combined visualization that color-coded each vertex for its loading in the two-dimensional PC space. To aid interpretation of the developmental trajectory variation captured by the resulting PCs, we modelled and visualized age effects on surface area for vertices residing at the extremes of the PC loading space (i.e., for two-dimensional model: high PC1/low PC2, low PC1/high PC2, high PC1/high PC2, low PC1/low PC2). Each of these four extremes was defined by taking the top 10% of vertices based on the sum of each vertex PC1 and PC2 loading, weighted by the desired extreme. For example, to find the extreme vertices for high PC1/low PC2, we computed PC1 loading * (−1*PC2 loading) for each vertex and took the top 10% of a ranked list based on this score.

## RESULTS

### Participant characteristics

Our longitudinal study included 1,529 structural magnetic resonance imaging scans from 792 healthy males and females (403 M/ 389 F) between 5 and 25 years of age (see **Table 1**).

### Total Amygdala and Hippocampal Volume Increase Over Development in a Curvilinear, Tri-phasic, and Sexually Dimorphic Manner

Visual inspection of raw volume data, and fitted trajectories for each structure, revealed large inter-individual variation in absolute structure size and rates of volume change, but a broadly tri-phasic growth pattern could be discerned across both structures and both sexes: robust volume increases during childhood, a phase of decelerating growth over adolescence, and gradual stabilization of bulk volume into early adulthood (**Fig 1**). Total bilateral hippocampal and amygdala volume were both significantly larger in males than females (p < 2*10^−16^) across the age-range studied. We also detected statistically-significant sex-differences in the shape of volume growth trajectories for the amygdala (p = 2.1*10^−7^) and hippocampus (p = 4.4*10^−8^).

**Figure 1.**
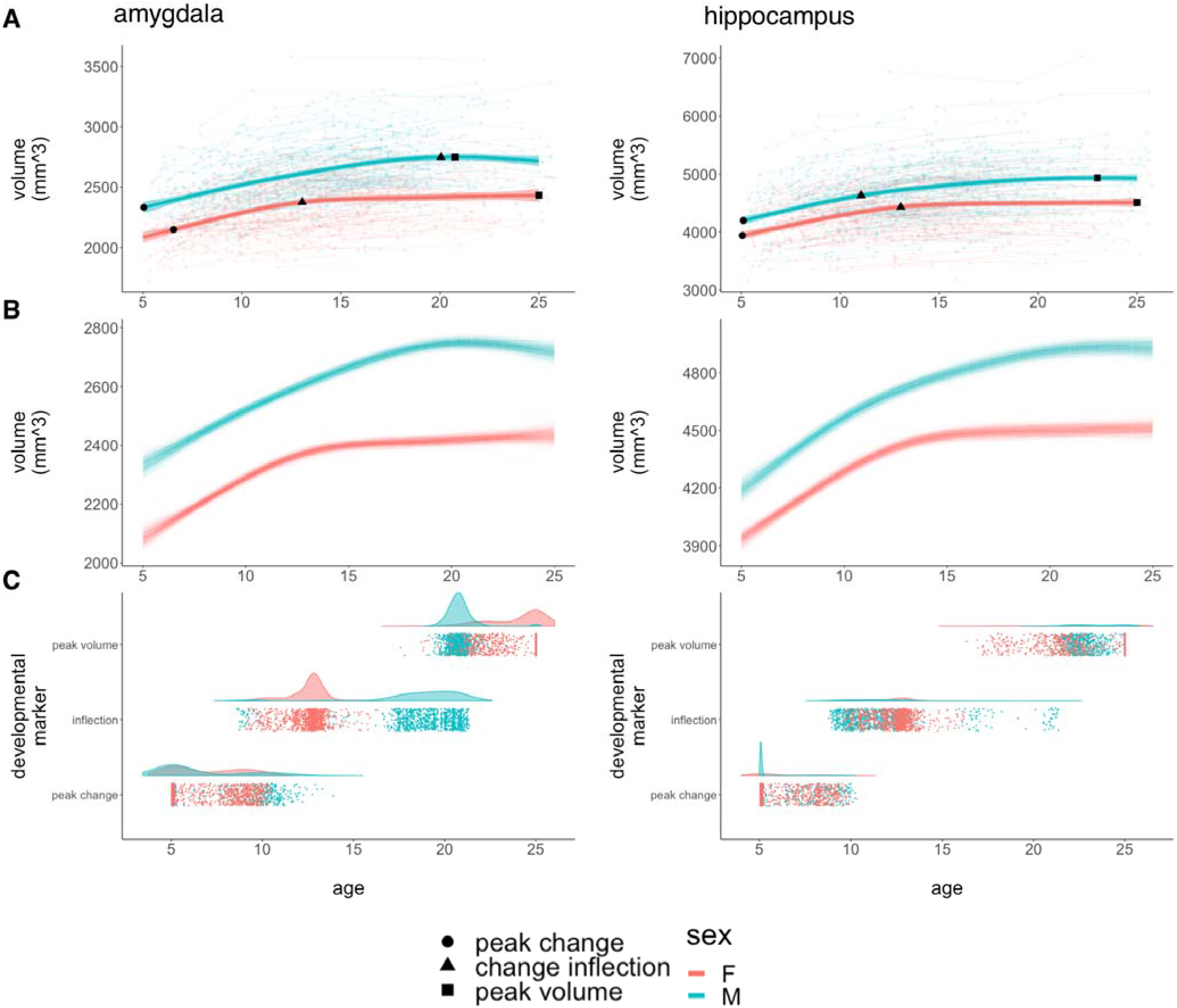
Developmental Curves and Milestones for Bulk Amygdalar and Hippocampal Volume. **A:** Spline-based, group-level trajectories for the bulk bilateral volume of each structure are shown by sex (males = blue / females = red). These fit lines are superimposed on spaghetti-plots of raw data showing individual volume measurements (background circles), linked by lines denoting observations from the same individual. Fit lines are surrounded by shaded 95% confidence intervals. The three developmental markers (i.e., age of fastest volume change [circles], age of greatest change in developmental tempo [triangles], and age at attainment of peak volume [squares]) overlay the fit lines for each sex and for each structure. **B:** Spline-based, group-level trajectories for the bulk bilateral volume of each structure from 1000 bootstrap samples of our data, where we resampled from the set of 792 individuals with replacement. **C:** Visualization of developmental milestone timing distributions across bootstrap samples, stratified by sex, where each point corresponds to the timing of a developmental milestone in a bootstrap sample.

Sex-differences in the trajectory of amygdala growth lead to a rapid accentuation of male-female differences in amygdala volume over mid-adolescence, with relative stabilization of volumetric sex-differences by the early 20’s (**Fig 1A**). This pattern was strikingly consistent across bootstrap resampling of our population (**Fig 1B**). Developmental milestone analysis further specified the key components of this sex-difference in amygdala growth (**Fig 1C, Table 2**): both sexes show comparably fast rates of amygdala volume growth during childhood, but females show a rapid deceleration in amygdala volume change at ~13 years, whereas amygdala growth does not undergo its most rapid deceleration in males until ~20 years of age (p_boot_=0.041). Notably, although this deceleration of amygdala growth occurs later and more gradually in males than females, males still achieve peak amygdala volume slightly earlier than females do (i.e., ~21 vs. 25 years), due to a continued gradual increase in female amygdala volume during the early 20’s that is absent in males (p_boot_=0.047).

**Table 2.** Age Estimates for Key Developmental Markers of Volume Change. Age of fastest volume change, age of greatest change in developmental tempo (i.e., inflection), and age at attainment of peak volume for the amygdala and hippocampus in each sex.

| | Male<br>Age<sub>empirical</sub> ($\mu_{boot}$ ; CI<sub>boot</sub>) | Female<br>Age<sub>empirical</sub> ($\mu_{boot}$; CI<sub>boot</sub>) | p<sub>boot</sub> |
| --- | --- | --- | --- |
| Age: Fastest Growth (95% CI)<br>Amygdala<br>Hippocampus | 5.1 (6.5 ; 5.1-11.1)<br>5.1 (5.7 ; 5.1-9.6) | 6.6 (6.9 ; 5.1-10.1)<br>5.1 (6.2 ; 5.1-9.5) | 0.329<br>0.436 |
| Age: Inflection (95% CI)<br>Amygdala<br>Hippocampus | 19.4 (18.8 ; 9.2-21.1)<br>11.0 (11.6 ; 9.2-20.9) | 12.8 (12.5 ; 9.8-14.4)<br>12.7 (12.4 ; 10.0-14.8) | 0.041<br>0.235 |
| Age: Peak Volume (95% CI)<br>Amygdala<br>Hippocampus | 20.8 (20.8 ; 19.8-22.2)<br>23.0 (23.4 ; 21.4-25.0) | 25.0 (24.0 ; 20.4-25.0)<br>25.0 (23.3 ; 18.0-25.0) | 0.047<br>0.389 |

Sex-differences in the trajectory of hippocampal growth were less overt than those for the amygdala (although still statistically significant at p = 4.4*10^−8^). These differences consisted of a more gradual accentuation of male-female differences in structure volume over the late teens due to faster volume increases in males relative to females (**Fig 1A, 1B**). Developmental milestone timing was more consistent between males and females in the hippocampus than in the amygdala (**Fig 1C, Table 2**). The developmental timing of volume growth, deceleration, and peak-attainment were broadly comparable between the amygdala and hippocampus (**Table 2**) – excepting the aforementioned male-specific delay in deceleration of amygdala growth.

### Spatially-Specific Sex-differences in Amygdala and Hippocampal Development

Sex-differences in the tempo of total amygdala and hippocampal volume were underpinned by highly focal, rather than spatially-diffuse, sex-differences in amygdala and hippocampal shape (**Fig 2**) as indexed by vertex-level measures of surface area. Specifically, vertex-level analysis detected “hot-spots” of sex-biased areal development (surviving FDR correction for multiple comparisons), which overlay the centromedial nucleus of the amygdala and extremes of the rostro-caudal hippocampal axis over CA1 and CA2. The sex-biased areal development observed at each of these foci (**Fig 2**) largely echoed the sex-biases observed for development of total amygdala and hippocampal size (**Fig 1**).

**Figure 2.**
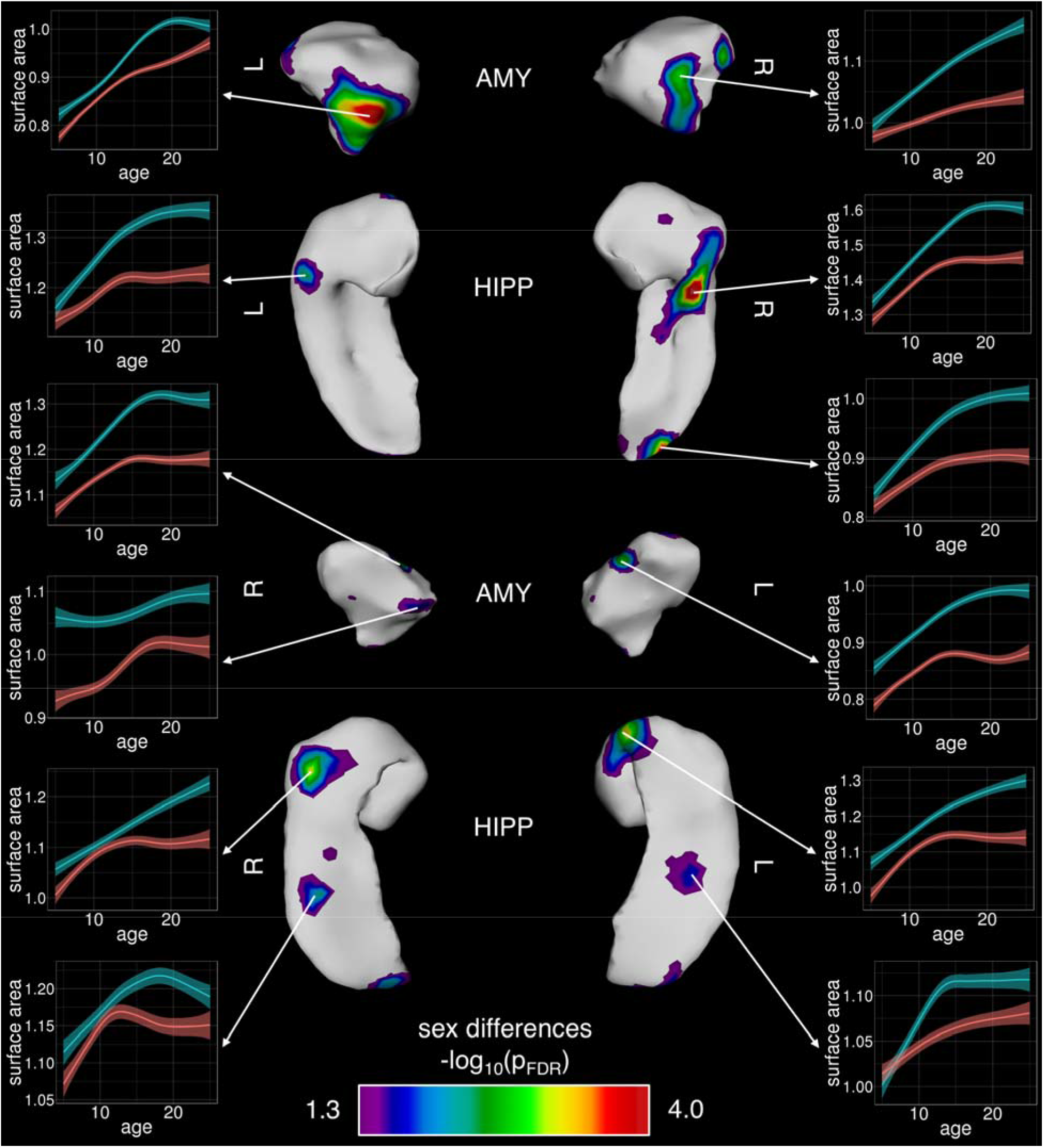
Foci of Statistically-Significant Sex-Differences in Development of Regional Amygdalar and Hippocampal Surface Area. Bilaterally symmetric areas of regional sex differences are shown across the left and right amygdala and hippocampus, where statistically-significant foci are illustrated post FDR-correction. Inset plots of surface area by age at individual vertices are provided to further illustrate male/female trajectory differences (males = blue/ females = red).

### Patterns of Anatomical Change are Topographically Organized Across the Amygdala and Hippocampus

We resolved spatial gradients of amygdalo-hippocampal change within each sex using two complementary approaches: vertex-wise mapping of percent area change, and a PCA of area change trajectories across all vertices. Percent area change over development showed a robust spatial gradient in both the amygdala and hippocampus. This broad gradient of developmental dynamism was reproducible in males and females, bilaterally symmetric, and coordinated between the amygdala and hippocampus (**Fig 3**). Regions of greatest area change within each sex coincided with regions of statistically significant differences in area change between sexes (cf. **Fig 2**), namely the centromedial nucleus of the amygdala and rostro-caudal extremes of the hippocampus. Regions of greatest area expansion were continuous across neighboring facets of the amygdala and underlying hippocampal head. Area change maps also revealed regions of the amygdala and hippocampus that are relatively stable in size across childhood and adolescence (**Fig 3**), most notably, the basolateral nucleus of the amygdala and caudal portions of the hippocampal tail (caudo-medial facets dorsally and caudo-lateral facets ventrally) spanning the subiculum and caudal extents of CA1.

**Figure 3.**
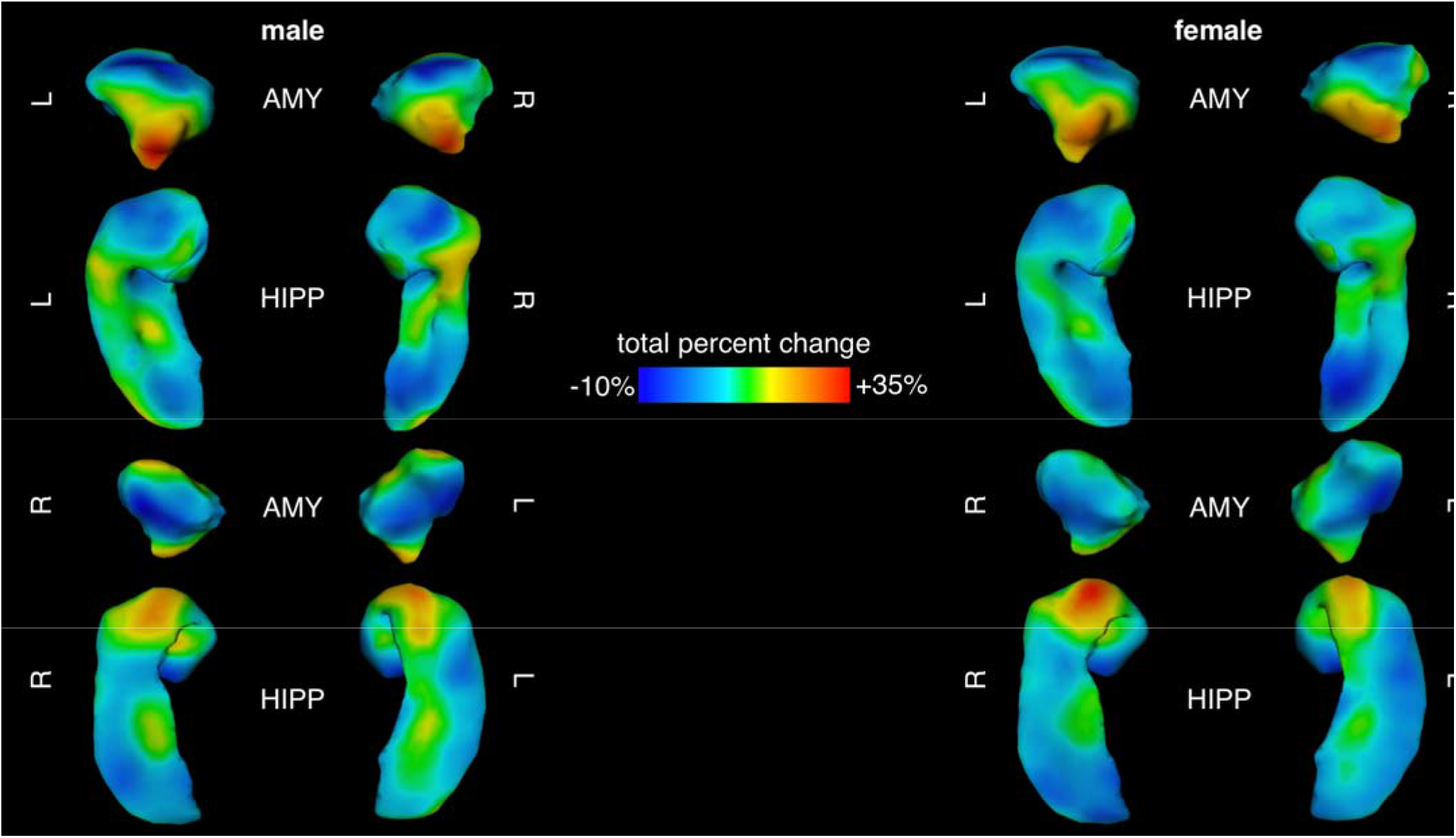
Percent Area Change Maps for Amygdala and Hippocampus Between 5 and 25 Years of Age in Males and Females. Maps of total percent change across the male (left) and female (right) amygdala and hippocampus, with blue representing the most stable areas (least dynamic) and red indicating areas of greatest areal expansion (most dynamic) from 5 to 25 years of age.

While these analyses identify clear spatial gradients in the magnitude of total area change across the amygdala and hippocampus, they cannot distinguish regional differences in the tempo of area change between ages 5 and 25 years. For example, a given vertex may increase in size by 20% through rapid acceleration in volume change over a few years with subsequent stasis, or through gradual increases that steadily accumulate across the whole age-range considered. To map tempos of amygdalo-hippocampal maturation in each sex, we modelled age-related changes in area for all 5,245 vertices across the amygdala and hippocampus, and then calculated first derivatives of these curves to provide vertex-specific estimates of annualized area change at 400 intervals between ages 5 and 25 years. Principal components analysis of this area change matrix in each sex (5,245 vertices by 400 age-point measure of annualized area change) identified 2 PCs that together explained >80% of the variance in the vertex-level area change trajectories (85.7% and 84.2% of variance in males and females, respectively). Vertex-wise PC loadings were also largely consistent between males and females - although more so for PC1 (Pearson’s *r* between sexes across vertices for PC1 loadings: 0.85) than PC2 (Pearson’s *r*: 0.50).

Visualizing both PCs separately in each sex (**Fig 4**) revealed that regional variation in PC1 loadings largely recapitulated those for total percent area change both qualitatively (c.f. **Fig 3**), and quantitatively (**Fig 5**; r: ~0.9 spatial correlation in vertex scores for area change and PC1 loadings in males and females). As for percent area change maps, the spatial patterning of PC1 was also similar between males and females (r=0.85 spatial correlation in vertex scores for PC1 loadings between males and females), and concordant for neighboring facets of the amygdala and hippocampus. Thus, the dominant dimension of amygdalo-hippocampal shape change between ages 5 and 25 years (variance explained in PC1: 69.5% males, 62.2% females) related to regional differences in overall magnitude of areal expansion achieved over development, which followed a similar topography in both sexes. In contrast to these findings for PC1, the spatial patterning of vertex loadings for PC2 **(Fig 4)** was distinct from that of total area change (**Fig 5**; r<0.4 spatial correlation in vertex scores for area change and PC2 loadings in males and females), and only moderately similar between males and females (**Fig 5**, r=0.51 spatial correlation in vertex scores for PC2 loadings between males and females). The proportion of trajectory variance accounted for by PC2 was also higher in females than males, in contrast to PC1, where the proportion of variance explained was higher in males versus females (variance explained in PC2: 16.2% in males, 22.0% in females).

**Figure 4.**
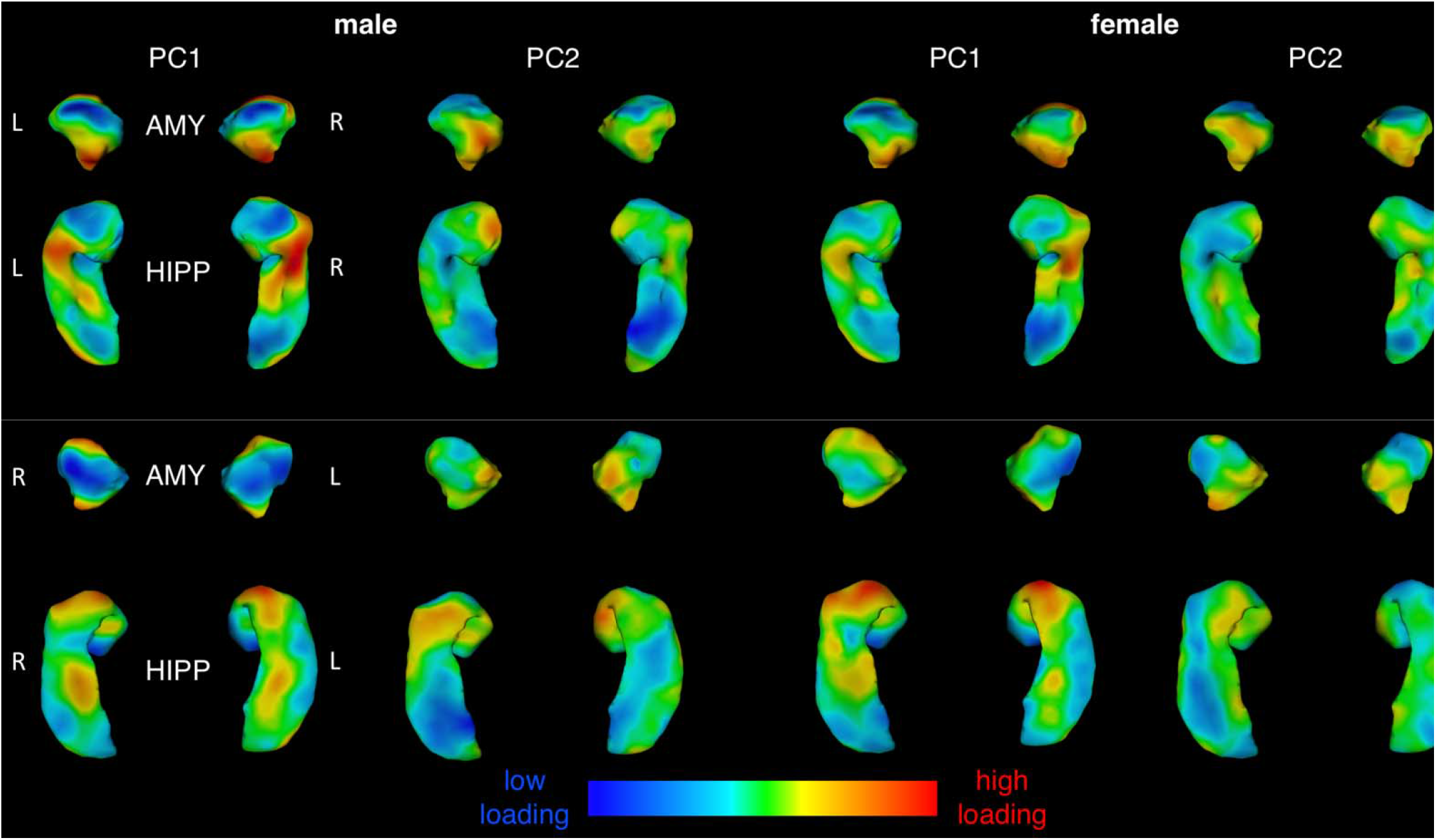
Surface Maps for Two Primary Principal Components of Amygdalo-Hippocampal Shape Change Between 5 and 25 Years of Age in Males and Females. Vertex loading maps for the two major components resulting from PCA of vertex-level amygdalo-hippocampal area change matrices, computed and shown separately for each sex.

**Figure 5.**
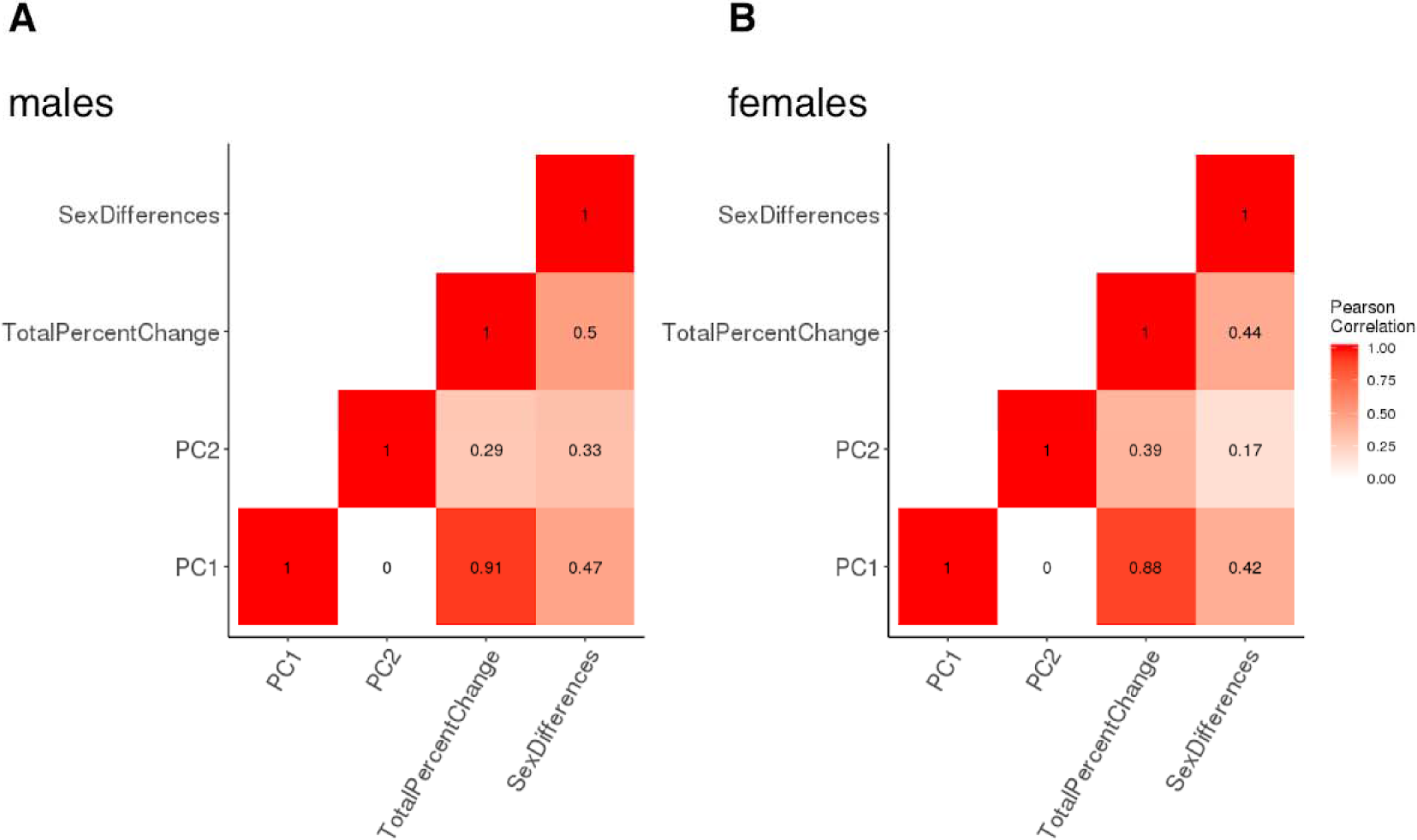
Heat Map of Spatial Correlations Between Different Surface Maps of Amygdalo-Hippocampal Area Change Between 5 and 25 Years of Age in Males and Females. Heat map visualization of correlations among PC1, PC2, total percent change, and sex differences. Darker colors indicate stronger correlations and lighter colors indicate weaker correlations. A separate heat map is provided for each sex.

To interpret PC2 while also taking into account the dominant gradient from PC1, we visualized both component loadings simultaneously in a color-coded, 2-dimensional space and plotted area trajectories for sets of vertices at all 4 extremes in the 2-dimensional PC loading space (**Fig 6**). In both males and females, variation in PC2 loadings had little discernable effect on the trajectory of area change amongst developmentally “static” regions with low PC1 loadings (i.e., red vs. green trajectories; **Fig 6**). However, amongst developmentally dynamic regions with high PC1 loadings - and more so in females than males - variation in PC2 loadings captured the distinction between regions with early/rapid growth deceleration vs. those with later/slower growth deceleration (purple vs. turquoise, respectively; **Fig 6**). This distinction showed its strongest topographic organization along the rostro-caudal axis of the dorsal amygdala in females, with developmentally-dynamic rostral vertices showing sharp growth deceleration in the early teens (purple), and developmentally-dynamic caudal vertices tending to show later and more gradual growth deceleration (turquoise). Males showed a similar rostro-caudal gradient in PC2 loadings within the amygdala, although this gradient captured less substantive regional differences in area change trajectories for males than females (in keeping with the lower proportion of variance explained by PC2 in males as compared to females). In both sexes, regional differences in PC2 loadings amongst developmentally-dynamic vertices showed weaker topographic organization in the hippocampus than in the amygdala (i.e., more spatially disjointed transitions from purple to turquoise; **Fig 6**).

**Figure 6.**
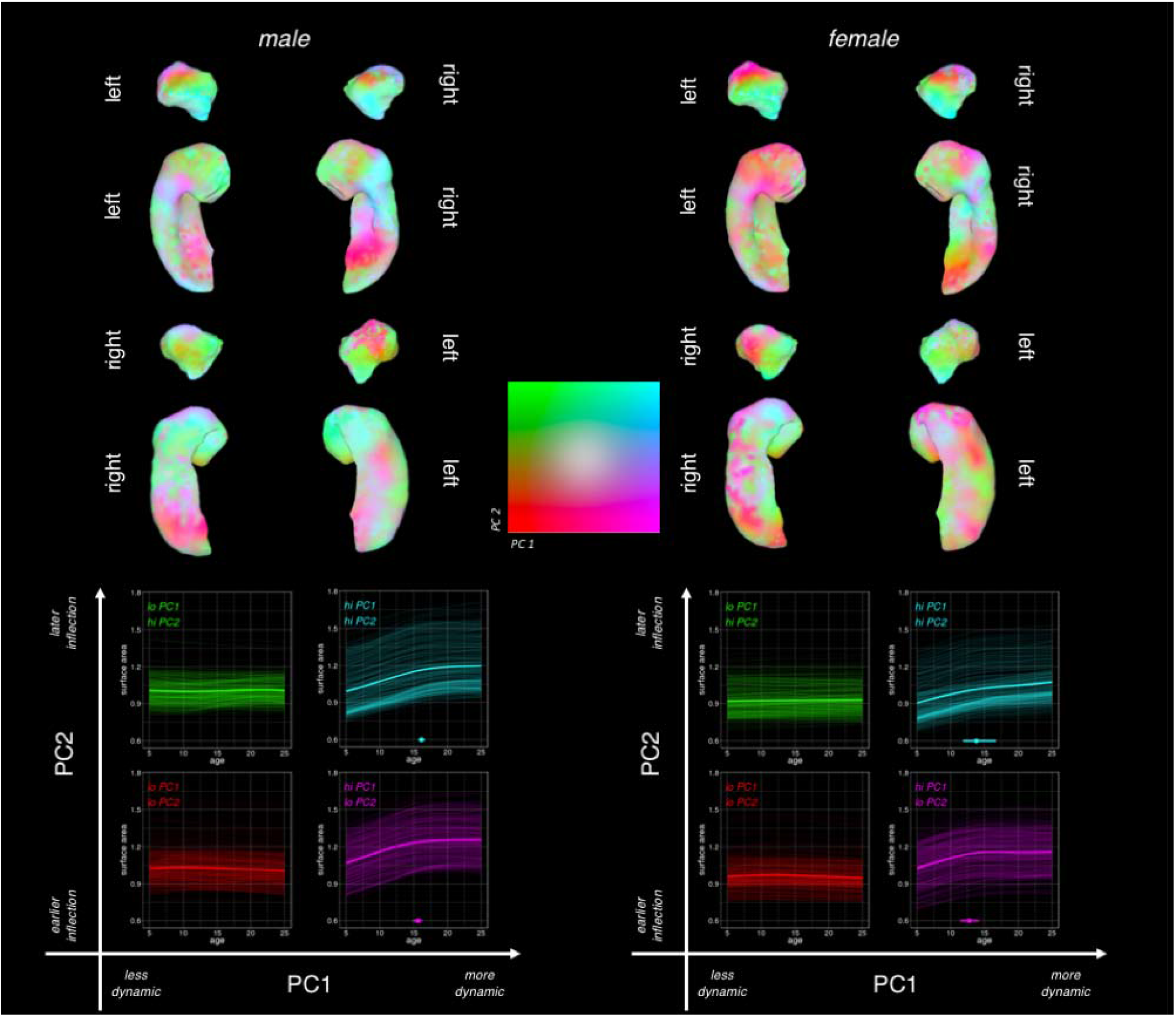
Combined Visualization for Both Principal Components of Amygdalo-Hippocampal Area Change Between 5 and 25 Years of Age in Males and Females. Simultaneous visualization of PC1 and PC2 across the left and right amygdala in both sexes separately, with PC1 loadings indicating level of dynamism (purple/ turquoise = more dynamic, green/ red = less dynamic), and PC2 loadings indicating inflection (green/ turquoise = early/ rapid growth deceleration, red/purple = later/slower deceleration). Bootstrap 95% confidence intervals are shown for the hi-PC1 quadrants of PC-space.

## DISCUSSION

### Development of Bulk Amygdala and Hippocampus Volumes

Our bulk volumetric analyses replicate past findings regarding sex differences in the absolute volume of both structures (males > females; Dennison et al., 2013; Herting et al., 2018; Nadig et al., 2018; Tamnes, et al., 2018; Wierenga et al., 2014). We also observed clearly non-linear trajectories of volume change for both structures which broadly replicate those detected in the other available longitudinal studies of amygdala and hippocampal volume trajectories across childhood and adolescence (Herting et al., 2018, Tamnes et al., 2018; Wierenga et al., 2014). Further replicating the largest of these earlier reports (despite our use of different image segmentation methods in an independent dataset) – we find significant sex-differences in the shape of amygdala and hippocampal growth trajectories. Collectively, these observations provide reproducible evidence for curvilinear and sex-biased trajectories of amygdalo-hippocampal volume change - using different cohorts, different image analysis tools, and different statistical models.

We also newly-define three key developmental markers to further characterize the timing and sex-differences in amygdalar and hippocampal volume development. For both structures, within the age-range considered, we observe the most rapid volume growth at approximately age 5-6 years in both sexes, followed by a phase of growth deceleration during the teens and a subsequent period of relative volume stability, with attainment of peak volume in males and females during the early 20’s. Our analyses isolate a ~7-year delay in the timing of growth deceleration for the amygdala in males vs. females (age 20 vs. 13 years, respectively), which may be relevant for the contemporaneous adolescent emergence of sex-biases in the prevalence of several psychiatric conditions in which the amygdala has been heavily implicated. These conditions include both female-biased mood/anxiety disorders (Price & Drevets, 2009), and the male-biased adolescent spike in behavioral dis-control and rule-breaking/conduct disorders (Jones et al., 2009). It is important to note, however, that different developmental milestones give different conclusions regarding sex-biases in the timing of amygdala maturation (e.g., attainment of peak amygdala volume is delayed in females relative to males), and resolving the behavioral relevance of milestone timing must await studies that can provide person-level estimations of milestone timing for multiple brain regions.

### Foci of Sex-Biased Amygdalo-hippocampal Development

Vertex-level analyses qualitatively localized sex-biased development to specific subregions of the amygdala (e.g., overlying centromedial nuclear groups) and hippocampus (e.g., overlying rostro-caudal extremes of the CA1 and CA2). Given that these amygdalo-hippocampal sub-regions are all thought to be important for affective processing (Kalin, Shelton, & Davidson, 2004; Plachti et al., 2019; Tillman et al., 2018), their sex-biased morphological development may be relevant for the well-documented sex-bias in risk for anxiety and mood disorders that emerges during adolescence (Rutter et al., 2003). Furthermore, the regionally-specific nature of these sex-biases in morphological development suggests a potential spatial targeting of sex hormone and/or sex chromosome influences on amygdala and hippocampal development in humans.

To date, most available data regarding mechanisms and microstructural markers for sex-biased amygdalo-hippocampal maturation have come from research in rodents (Corre et al., 2016; Hines, Allen, & Gorski, 1992; Qiu et al., 2018; Vousden et al., 2018). Most notably, rodents show a well-established stereotyped sex difference in volume of the medial nucleus of the amygdala (Hines et al., 1992) - which is larger in males than females throughout the lifespan. A recent longitudinal neuroimaging study identified a sharp increase in magnitude of this sex-bias during murine puberty (Qiu et al., 2018). The location (overlying medial amygdala), direction (male>female) and developmental timing (pubertal accentuation) of this anatomical sex bias in mice are all fully concordant with the findings we present here in humans – suggesting that humans may also share the same mechanistic basis for these macroanatomical sex-differences as mice. To date, existing experimental data from rodents suggest that sex biases in medial amygdala volume are first established in perinatal life through the action of circulating androgens and may also be maintained by ongoing actions of sex-steroids across puberty (De Lorme, Schulz, Salas-Ramirez, & Sisk, 2012; Morris, Jordan, & Breedlove, 2008). Murine neuroimaging data also suggest that medial amygdala volume differs as a function of chromosome complement in a manner that opposes the directional effects of gonadal sex-steroids (i.e., XY < XX, and gonadal males > gonadal females). Comparable experimental data are lacking in humans, but there is observational neuroimaging evidence for both sex steroid (Peper, Hulshoff Pol, Crone, & van Honk, 2011) and sex chromosome effects (Nadig et al., 2018) on medial amygdala organization in humans.

Although the hippocampus is not a classical focus of sex-biased brain volume in mice, recent neuroimaging studies have identified a dorsal hippocampal region where volume is greater in male vs. female mice (Qiu et al., 2018). This region is considered to be comparable to the extreme caudal hippocampal tail in humans (Strange et al., 2014), which we also find to be larger in males than females. As for the amygdala, observational neuroimaging data suggest that human hippocampal volume may also be sensitive to sex steroid (Peper et al., 2011) and sex chromosome (Nadig et al., 2018; Satterthwaite et al., 2014) effects. Thus, existent data – though scant – are consistent with the idea that there are conserved sex-biases in regional amygdalo-hippocampal organization, and that these differences may arise through the combined effects of sex-differences in chromosomal complement and gonadal status.

### Spatial Gradients of Amygdalo-hippocampal Maturation

Our findings complement published analyses of subnuclear volume maturation (Tamnes et al., 2018) by providing the first spatially fine-grained maps of shape change within the amygdala and hippocampus across human childhood, adolescence, and early adulthood. These vertex-level maps suggest that amygdalo-hippocampal maturational gradients vary both within and across classical subnuclear boundaries. The independent recovery of the topographical gradients in males and females effectively provides split-half evidence for reproducibility of these two principal axes of amygdala-hippocampal maturation in humans.

The first principal component of amygdalo-hippocampal shape change (PC1) between childhood and early adulthood captures regional variation in the total magnitude of surface area expansion over development. High PC1 loadings are seen at rostral and caudal extremes of the amygdala (separated by a relatively developmentally-static central domain), and neighboring facets of the hippocampal head, as well as a more caudally-located region spanning the lateral hippocampal “neck” and medial mid-hippocampal tail. Most foci of sex-biased shape development (**Fig 2**) lay within regions of high PC1 loading (**Fig 4**). Amongst regions with high PC1 loadings, the second principal component of amygdalo-hippocampal shape change (PC2) captured variation in the timing of growth deceleration.

In the amygdala, and especially so in females, PC2 distinguished regions overlying the basolateral nuclei (developmentally dynamic, with relatively fast and early growth deceleration) from those overlying the centromedial nucleus (developmentally dynamic with later and slower growth deceleration). Thus, developmental tempos of anatomical expansion within the amygdala may be organized by the distinct functional and connectivity profiles of different amygdala subnuclei (Sah et al., 2003; Swanson & Petrovich, 1998). In contrast, the spatial patterning of PC1 and PC2 within the hippocampus bore little relationship with the spatial distribution of different hippocampal subfields. However, there is growing evidence that hippocampal functional specialization is also organized along a rostro-caudal axis that runs orthogonal to the orientation of cytologically-defined hippocampal subfields (Brunec et al., 2018; Collin, Milivojevic, & Doeller, 2015; Plachti et al., 2019; Poppenk et al., 2013; Sah et al., 2003; Vos de Wael et al., 2018). Indeed, there are notable conjunctions between our map of hippocampal anatomy change with age and a recent parcellation of the hippocampus based on its functional connectivity with other brain regions (Plachti et al., 2019). For example, connectivity-based parcellation defines a segment of the medial hippocampal tail bilaterally which is preferentially activated during autobiographical memory tasks, and notable for showing little anatomical change over development in our study (**Fig 3**). We also observe convergence between connectivity-based parcellation and our developmental maps over latero-rostral facets of the hippocampal tail. These regions - which are notable for their highly dynamic and sex-biased areal expansion over development in our analyses - also emerge as distinct clusters in connectivity-based parcellation of the hippocampus, with tendency for preferential activation during emotion processing. Thus, the developmental changes in hippocampal shape may be more closely organized by functional rather than cytological territories.

### Limitations

Our findings should be recognized in light of certain limitations. First, all scans were acquired on a 1.5 Tesla scanner, and future large longitudinal studies at higher field strengths (Casey et al., 2018) will be able to improve the signal-to-noise ratio and detection of the boundaries that occur around and within each structure. Ultra-high-field imaging – although less well suited for longitudinal applications in children – will be helpful for assessing how foci of sex-biased shape development are related to the underlying microstructural features that define classical amygdalo-hippocampal subnuclei/subfields. Second, our image analysis methods are focused on volume and shape as anatomical phenotypes, and simultaneous consideration of multiple in-vivo imaging phenotypes (Seidlitz et al., 2018) will be an important next step in refining our understanding of amygdalo-hippocampal development in humans. Third, our emphasis on trajectory mapping between childhood and early adulthood required us to collapse across Tanner stages in order to maximize the number of data points across our full age-range of interest. It is likely, however, that chronological age and pubertal status interact in complex, non-linear ways to influence brain maturation (Forbes & Dahl, 2010; Goddings et al., 2014; Herting & Sowell, 2017; Satterthwaite et al., 2014; Wierenga et al., 2018). Future studies in larger samples will be required to untangle age and pubertal effects on non-linear anatomical growth, and to tackle the broader question about how to best measure pubertal progress. Larger datasets, with more complete longitudinal data will also be required to test if inter-individual variation in trajectories of amygdalo-hippocampal maturation can be related to trajectories of cognitive/behavioral change, and if any such relationships differ between males and females.

Despite these limitations, our study represents the largest longitudinal analysis of amygdalo-hippocampal development in health within a single-site cohort to date. These results provide new insights into the spatiotemporal patterning of amygdalo-hippocampal development, highlighting an integrated architecture and focal sex-differences that may contribute to adolescent-emergent male-female differences in behavior and psychopathology.

## ACKNOWLEDGMENTS

The authors would like to thank the participants and their families for their participation in this study. AN and CLM are supported by NIH Intramural Research Training Awards. PKR and JS are supported by the NIH Oxford-Cambridge Scholars’ Program. This work was supported by the intramural program of the National Institutes of Health (Clinical trial reg. no. NCT00001246, clinicaltrials.gov; NIH Annual Report Number, ZIA MH002949-03).

